# The effect of quercetin on the fluorescent intensity of neurofibrillary tangles that correspond to Alzheimer’s disease within *Drosophila melanogaster*

**DOI:** 10.1101/2025.09.20.677467

**Authors:** Gavin Carey, Ananya Nadooli

**Affiliations:** Academies of Loudoun

## Abstract

Alzheimer’s disease (AD) is a neurodegenerative disease that accounts for more than 60% of all dementia cases, and over seven million Americans above 65 years old were affected with AD in 2024 (Gaugler et al., 2024). A symptom of AD is severe memory loss, which leads to lowered brain function and a need for intensive care. Currently, AD only has five symptomatic approved drugs. However, there is a lack of research in the progression of AD through the neurofibrillary tangle mechanism. Tau is a microtubule stabilizing protein that becomes unstable when hyperphosphorylated. Hyperphosphorylated tau misfolds, resulting in toxic aggregates that accumulate to create neurofibrillary tangles. Saffron decreased tau fibrillation, neurotoxicity and slowed the neurofibrillary tangle formation (Patel et al., 2024). Therefore, quercetin, an antioxidant present in saffron, was hypothesized to reduce the intensity of the neurofibrillary tangles within *Drosophila melanogaster*, commonly known as the fruit fly, exhibiting AD. *Drosophila melanogaster* has easily modifiable genes and shares a similar genetic sequence with humans, which is ideal to research if the effect of a supplement in flies could have a similar effect on humans. This study found that quercetin had no significant effect on the intensity of the neurofibrillary tangles within *Drosophila melanogaster*. The ANOVA statistical test showcased that the p-value was above 0.05 between flies exhibiting AD and flies exhibiting AD with supplemented quercetin. Possible error sources include the fluorescence of brain tissue present and the large constant area, (35 u^2^), measured for each brain image.

## Introduction

Alzheimer’s disease is a neurodegenerative disease that causes dementia and early death within elderly populations. Around ten percent of Americans that are 65 years or older are suffering from some form of dementia, with Alzheimer’s disease being the most common cause (Manly, 2022). Alzheimer’s disease itself is the sixth leading cause of dementia in the United States, and as of 2025, there is currently no cure for the disease. Dementia caused by Alzheimer’s disease results in difficulties in cognitive function and a decline in motor skills. Loss of memory is due to the neurodegeneration of the brain, which is the death of millions of neurons. Additionally, neurons do not come back because the neurons will never enter mitosis to split and produce new cells once the organism has a certain amount. This causes the death of these neurons to have drastic and detrimental effects, as the damage done to these neurons is irreversible since the neurons can not be replaced. There are three main mechanisms that lead to neurodegeneration within the brain: Aβ plaque aggregation, oxidative stress, and aggregation of hyperphosphorylated tau into neurofibrillary tangles. Neurofibrillary tangles are accumulations of tau proteins, which are microtubule-associated proteins found within the neurons (Khan et al., 2020).

Aβ plaques, derived from the breakdown of a protein called amyloid precursor, result in neurodegeneration through inflammation within the brain that activates the microglia, also known as the immune cells of the brain. The activation of microglia eventually leads to neuronal death in the brain. The constant activation of the microglia within the brain makes removing the plaques from the brain difficult. In response to the accumulation of Aβ, the microglia activate and produce proinflammatory cytokines, which leads to a permanent state of neuroinflammation. This state of neuroinflammation worsens neurodegeneration and further progresses Alzheimer’s disease (Miao et al., 2023). Oxidative stress, or the imbalance of free radicals and antioxidants in the body, is also present in the brain when leading to neuronal death and neurodegeneration (Pizzino et al., 2017). Some products of oxidative stress include neurofibrillary tangles and Aβ plaques. Oxidative stress can cause oxidative change, the chemical change that uses the transferring of oxygen among atoms, within the Aβ plaques making them more toxic and making their removal harder (Khan et al., 2020).

A major mechanism of Alzheimer’s disease progression is neurofibrillary tangles made of hyperphosphorylated tau, a protein found in the brain that binds to microtubules. Microtubules are components of the cytoskeleton, and the tau maintains the integrity of brain cells when bound to microtubules (Lasser et al., 2018). Tau can become unstable when hyperphosphorylated, which leads to the dissociation of tau from microtubules. The dissociation of tau from the microtubules causes the tau to misfold, and this misfolding leads to an increase in tau to tau interactions. As a result, the interaction of tau produces the formation of toxic aggregates, known as neurofibrillary tangles (Zhang et al., 2021). Tau misfolding leads to the inability of tau to bind to microtubules, which causes the microtubules to be unable to uphold the structural integrity of the cells (Khan et al., 2020). Tau pathology spreads among cells and causes the misfolding of more tau proteins, which create neurofibrillary tangles. Misfolded tau proteins act as seeds that spread, creating toxic accumulations of tau within the form of neurofibrillary tangles (Zhang et al., 2021). Further accumulation of tau aggregates facilitates the spread of tau pathologies throughout the brain and results in tau toxicity that leads to neuronal death and the further onset of dementia. Tau spreads through anatomically connected regions, so neurofibrillary tangles originate in the hippocampus and then later spread throughout the brain, influencing the degeneration of neurons in all parts of the brain (DeTure & Dickson, 2019). Therefore, tau aggregation is considered the major mechanism influencing neurodegeneration within Alzheimer’s disease (Trejo-Lopez et al., 2022) The most efficient way to slow the progression of tau pathologies is by binding the tau before the protein becomes misfolded, thereby preventing the tau from forming toxic aggregates by ensuring that the tau maintains the original shape (Kashif et al. 2024).

There are many models that are used to study neurodegenerative diseases. *Drosophila melanogaster*, commonly known as the fruit fly, is the model that was used during experimentation and is one of the most popular models for studying neurodegenerative diseases. *Drosophila* are low cost model organisms with a short lifespan, making fruit flies an ideal organism for studying neurodegenerative diseases. Additionally, the brains of *Drosophila melanogaster* are compact, which is beneficial for the study of neurodegenerative diseases (Nitta & Sugie, 2022). The genetics of fruit flies can easily be altered, which helps researchers to express diseases related to genetics and allow for the expression of proteins. For example, Alzheimer’s disease can be researched and expressed in *Drosophila melanogaster* through the expression of tau in the neurons of the brain. Changes to the DNA of the *Drosophila melanogaster* can be made through the use of the GAL4 and UAS system, in order to add and activate genes that create proteins within certain areas of the *Drosophila*. The GAL4 line is the yeast transcriptional factor, and the GAL4 line provides the area where the UAS will activate and produce a protein. UAS stands for upstream activation system, and the UAS expresses a protein in the part of the body that is chosen by the GAL4 system by activating specific genes that lead to the production of the chosen proteins. The protein that will be expressed will be human tau, and the human tau will be expressed in the neurons. C155 Elav is the name for the GAL4 system that is associated with the area of the brain, which will be used during the project. The protein, tau, is only expressed when both the GAL4 and the UAS are present in the same fly, and the expression is achieved through the crossing of the flies’ genetic lines through breeding. Flies with the genes of elav-GAL4/CyO and UAS-MAPT are crossed to produce progeny exhibiting Alzheimer’s disease. The parent genotypes would be yw; UAS-MAPT and w^-^; elav-GAL4/ CyO. The elav-GAL4 and CyO will be the male *Drosophila melanogaster* genes and the UAS-MAPT will be the female *Drosophila melanogaster* genes. Progeny containing elav-GAL4 and UAS-MAPT will exhibit Alzheimer’s disease. Progeny containing the UAS-MAPT and CyO genes will have curly wings. Around half of the progeny will have curly wings and the other half will exhibit Alzheimer’s disease. The progeny we will be looking for is elav-GAL4/UAS-MAPT, and we will distinguish between the two different progeny by looking for the progeny without curly wings. Since these two combinations are the only possible gene combinations present in the progeny, the progeny without curly wings will have Alzheimer’s disease and exhibit human tau within them. Therefore, flies without curly wings will be used for experimentation. A punnett square of the cross of the GAL4 line and the UAS line will be shown below in Figure 1.

**Figure 1:**
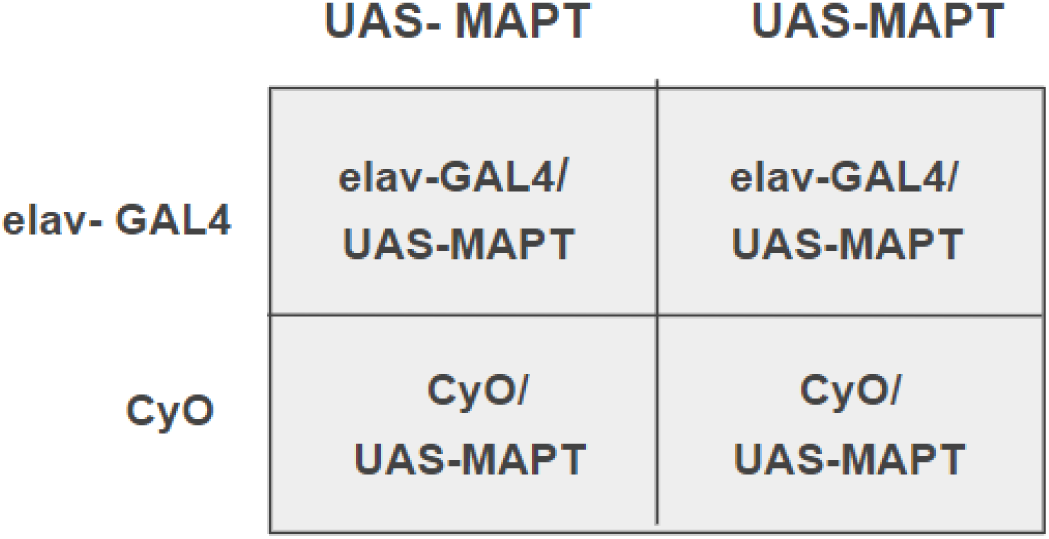
Punnett Square for the Genetic Cross (**elav-GAL4/CyO × UAS-MAPT**)

A main way to reduce neurodegeneration during the progression of Alzheimer’s disease is to consume foods and substances that are filled with antioxidants. One such substance is the common herb, saffron. Saffron has many key antioxidants and vitamins that cause the herb to have neuroprotective effects, such as slowing the progression of Alzheimer’s disease and attenuating memory loss. One of the main antioxidants that is found in saffron is called quercetin (Avila-Sosa et al., 2020). The neuroprotective effects of saffron could be attributed to saffron’s high antioxidant and anti-inflammatory properties. In the past, saffron has been researched to impact neurofibrillary tangles, a key neuropathological component of the progression of Alzheimer’s disease. Saffron attenuated the buildup of neurofibrillary tangles and Aβ plaques within rats, as well as decreasing neurodegeneration, leading to the saffron group having a milder exhibition of Alzheimer’s in comparison to the control group. The introduction of saffron into the diet of rats was connected to decreased neurodegeneration, cognitive impairment, and memory loss when compared with the control group of rats unsupplemented with saffron that exhibited Alzheimer’s disease (Patel et al., 2024).

Quercetin is a flavonoid, a specific type of antioxidant that has potent effects on health through its neuroprotective and antioxidant properties. Flavonoids are phytochemical compounds that are present in many different natural substances such as fruits, herbs and seeds. Flavonoids have antitumor, antioxidant, and neuroprotective properties (Ullah et al., 2020). Quercetin reduces the accumulation and toxicity of Aβ plaques within the Alzheimer’s disease brain by increasing the activity of the enzymes that degrade Aβ. Quercetin also has benefits for heart diseases, as quercetin lowers blood pressure in overweight and obese adults (Carrillo-Martinez et al., 2024). Flavonoids are also used as antiviral and antibacterial compounds. Flavonoids obstruct the entrance of viruses into cells and inhibit the viruses from being released to spread to other cells, spreading the virus throughout the body. Flavonoid compounds also can disrupt the replication process of viruses, making the spread of the virus throughout the body difficult (Badshah et al., 2021). Moreover, this is just one of the many medicinal uses of flavonoids, which displays its potential and versatility in future research.

Quercetin has potent binding properties, which could have implications on the ability to to prevent the progression of neurofibrillary tangles by binding tau and to display strong neuroprotective factors, such as the lowering the oxidative stress and anti-inflammatory properties within the brain (Carrillo-Martinez et al., 2024). Quercetin has the ability to bind transthyretin, a protein that loses its structure leading to fibril formation like tau. Quercetin has the ability to stabilize the protein, preventing the transthyretin from forming fibril tangles (Ciccone et al., 2024).

Another antioxidant that had the ability to bind transthyretin was curcumin, demonstrating that this antioxidant has similar binding abilities to quercetin as they are both able to bind the same thing and prevent transthyretin from forming these tangles. Curcumin, also found in saffron, is another antioxidant that showcased strong binding abilities and anti-inflammatory properties (Goel & Aggarwal, 2010). The binding of transthyretin by curcumin prevented the formation of fibril tangles in mice and also inhibited further aggregation. Curcumin also lowered the toxicity of the transthyretin fibrils (Ferreira et al., 2019). Curcumin demonstrated a high binding affinity for tau, the protein that forms neurofibrillary tangles. The high binding affinity impeded the aggregation of tau, slowing neurodegeneration and the progression of Alzheimer’s disease. Curcumin consumption also enhanced cognitive function in patients exhibiting Alzheimer’s disease (Kashif et al., 2024).

The binding of tau, which prevents the formation of neurofibrillary tangles, has a strong effect on the neurodegeneration and the number of neurofibrillary tangles within the brain. An antibody named semorinemab demonstrated the protection against tau toxicity and attenuated tau pathology, which was achieved by binding the tau prior. The antibody achieved protection against tau toxicity and attenuated tau pathology by binding the tau before the protein aggregated and had turned into toxic neurofibrillary tangles (Ayalon et al., 2021). The binding of tau was shown to decrease the accumulation of tau and the number of tau pathologies within the brain of the mice tested on. Furthermore, the binding of the tau protein prevents the protein from misfolding and turning into the toxic aggregates that spread throughout the neurons, which protects the neurons by preventing neurodegeneration (Mroczko et al., 2019). Thus, through the prevention of misfolding and the protection of neurons, the binding of tau could impair the progression of Alzheimer’s disease.

However, with the light of new possibilities for the prevention and treatment of Alzheimer’s disease research, there have been many problems that have arisen in this field of study as well. Overall, Alzheimer’s disease has only five “symptomatic” approved drugs, which has shown to be a challenge in drug development for Alzheimer’s disease (Luca, 2018). Moreover, ten or more years might be needed in order for clinically tried drugs to be brought to the market, which has been an issue in the field of Alzheimer’s disease research since the long testing period hinders the ability to produce personalized drugs with greater efficiency for those who struggle with Alzheimer’s disease (Veurink et al., 2020). Furthermore, a healthy lifestyle through beneficial diets, such as those with antioxidants, was shifted to be a more holistic approach to research. However, while current research has shown that antioxidants play a role in reducing the progression of Alzheimer’s disease, there have been issues in the field to provide strong clinical evidence for this claim, and there has been a lack of research in the progression of Alzheimer’s disease through neurofibrillary tangles (Singh & Ghosh, 2018).

The purpose of this study is to determine the effect that quercetin has as an oral supplement on the neurofibrillary tangles modeled by *D. melanogaster* that exhibit Alzheimer’s disease. Specifically, this study aims to determine whether there is a correlation between the amount of quercetin taken as a dietary supplement and the intensity of neurofibrillary tangles. Research into this subject will display whether quercetin has the potential to be an agent in treating Alzheimer’s disease by reducing the amount of neurofibrillary tangles within the brain. The hypothesis was that if *D. melanogaster* consume quercetin, via their daily food intake, then the intensity and number of neurofibrillary tangles within their brain will decrease due to quercetin’s ability to bind tau. To test the correlation, quercetin will be supplemented within the food of the fruit flies. The experimental group of *D. melanogaster* that exhibit Alzheimer’s disease and are supplemented with quercetin will be compared to four different control groups. The negative control was *Drosophila melanogaster* from either the GAL4 that does not intake quercetin. The positive control was *Drosophila melanogaster* that exhibit Alzheimer’s disease but do not intake quercetin in their diet. The toxicity control was flies from the GAL4 line that intake quercetin. The negative control helped in order to study the flies that do exhibit Alzheimer’s disease, as the neurofibrillary tangles in the brain between flies that do and do not undergo genetic modification can be compared. The negative control allowed us to analyze the effects of the treatment on normal patients to see if there are any adverse effects. This toxicity control would present data on fruit flies that do not have Alzheimer’s disease. The treatment toxicity allowed for the comparison of the neurofibrillary tangles in relation to quercetin without the flies undergoing genetic modification to exhibit Alzheimer’s disease. Moreover, this group will be used to analyze the effect that quercetin has on *Drosophila melanogaster*. The positive control was *Drosophila melanogaster* that exhibits Alzheimer’s disease but doesn’t intake quercetin, which would allow for measuring the effect of quercetin. The comparison between flies that do not intake quercetin and those that do intake quercetin will be used to analyze the effect that quercetin has on the neurofibrillary tangles in *D. melanogaster* that exhibit Alzheimer’s disease.

The two independent variables within this experiment are the presence of Alzheimer’s disease in the *D. melanogaster* and the presence of quercetin in the daily food intake of the flies. A toxicity assay, which tests the effect of different concentrations on an organism’s lifespan, was performed to analyze the maximum amount of quercetin that the fruit flies can safely consume. Two concentrations were chosen based on the data received from this assay. Furthermore, the dependent variable of this experiment is the fluorescent intensity of the neurofibrillary tangles within the brain of the *Drosophila melanogaster*, which will be measured using a fluorescence microscope after the brain is stained with Thiazine Red. *D. melanogaster* that exhibit Alzheimer’s disease will be supplemented with quercetin through their daily food intake, and after a two week period, the brain of *D. melanogaster* will be stained using Thiazine Red. Thiazine Red is histochemical that is a marker of dense plaques and aggregates (Kniewallner et al., 2016). Thiazine Red binds itself to the tau that makes up neurofibrillary tangles after the brain is stained, causing the neurofibrillary tangles to show up on the fluorescence microscope and glow a bright red color. A lower fluorescent intensity correlates to a lower amount of tau in the brain. Since neurofibrillary tangles are accumulations of tau proteins, a lower amount of tau shows that there is a lower amount of neurofibrillary tangles within the brain. If the color of the brain sample is intense under the fluorescence microscope, then there is a large number of neurofibrillary tangles within the sample. The number of plaques that showed up under the microscope after the plaques were stained had a high correlation with the number found within the brain through other methods (Kniewallner et al., 2016).

Research done on this subject will provide further insight into the mechanisms of Alzheimer’s disease, and the research could show a possible path for a treatment of Alzheimer’s disease. This study would provide insight into the ability of antioxidants, specifically quercetin, to bind with the tau inside the brain. Other antioxidants like quercetin, such as curcumin, have previously been shown to impede the aggregation of tau. Moreover, the common herb saffron, which contains many antioxidants, including quercetin, had also displayed strong binding abilities and the prevention of neurofibrillary tangles. However, the effect of quercetin itself has never been studied on tau and neurofibrillary tangles. This research would help to take a critical step in Alzheimer’s disease treatments and prevention, as there is currently no cure for Alzheimer’s disease. If quercetin decreases the number of neurofibrillary tangles within the brain and improves the state of the *Drosophila melanogaster* suffering from Alzheimer’s disease, then other antioxidants could be researched to find a potent way in order to prevent the progression of the disease and to stop individuals from getting the disease altogether. Additionally, this result would indicate that neurofibrillary tangles are a major driver of the progression of Alzheimer’s disease, which allows researchers to determine what to target in order to either cure or stop the progression of Alzheimer’s disease. Many studies have been done on antioxidants on Alzheimer’s disease as oxidative stress is a key component of the progression of Alzheimer’s disease, but this approach in reducing the progression of Alzheimer’s disease has not been considerably explored. The effect of quercetin as an oral supplement on neurofibrillary tangles has never been studied before. Furthermore, quercetin as the specific antioxidant to reduce the neurofibrillary tangles in the brain has never been studied as well. Furthermore, data found on the flies could potentially translate to having a human impact, since *Drosophila melanogaster* and humans have a similar genetic makeup. Possible findings could potentially have an impact on Alzheimer’s treatment and prevention in humans, which would improve the lives of patients by preserving their memory and lengthening their projected lifespan. Research done on the use of antioxidants and binding of tau could have massive ramifications on the further study of Alzheimer’s disease by providing a direction for further research in order to prevent and to possibly find a cure for this life-altering disease. Given the prominence of Alzheimer’s disease, research into this topic is further encouraged, as these results could take a crucial initiative in treating millions of Alzheimer’s patients around the world.

The constants of this experiment are the lab equipment, the location at Dr. Eliason’s room, the time of day from 9-11 AM EST, the age of 3 weeks for the flies, the temperature of 22°C, and the female sex of the flies that are tested. There were three control groups for this experiment: one negative control, one toxicity control, and one positive control. The negative control was *Drosophila melanogaster* from the GAL4 line that does not intake quercetin. The positive control was *Drosophila melanogaster* that exhibits Alzheimer’s disease but does not intake quercetin, and the toxicity control was flies from the GAL4 line that intake quercetin. The line used for the controls stayed constant throughout the experimentation, and so the GAL4 line was used for all controls. The negative control helped in order to study the flies that do exhibit Alzheimer’s disease and don’t intake quercetin, as the neurofibrillary tangles in the brain between flies that have and have not undergone genetic modification can be compared. Moreover, the negative control also allowed us to analyze the effects of the treatment on normal patients to see if there are any adverse effects. This toxicity control presented data on fruit flies that do not have Alzheimer’s disease. The toxicity control allowed for the comparison of the neurofibrillary tangles in relation to quercetin without the flies undergoing genetic modification to exhibit Alzheimer’s disease. Moreover, this group was used to analyze the effect that quercetin has on *Drosophila melanogaster*. The positive control was *Drosophila melanogaster* that exhibits Alzheimer’s disease but doesn’t intake quercetin, which allowed for measuring the effect of quercetin. The comparison between no intake of quercetin and the intake of quercetin on Alzheimer’s disease flies was used to analyze the effect that quercetin has on the neurofibrillary tangles in *D. melanogaster* that exhibit Alzheimer’s disease.

The expected outcomes for fluorescent intensity with our groups from lowest to highest were the toxicity control group, the negative control, the experimental group, and the positive control group, respectively. The groups and expected levels of fluorescent intensity will be shown below in figure 2. A lower fluorescence intensity corresponds to a lower amount of neurofibrillary tangles, and a higher fluorescent intensity corresponds to a higher amount of neurofibrillary tangles. If quercetin is able to decrease the aggregation of tau into neurofibrillary tangles as hypothesized, and Alzheimer’s disease is associated with a high amount of neurofibrillary tangles, then flies with Alzheimer’s disease would have a higher fluorescent intensity than those without and flies that are not supplemented with quercetin should have higher fluorescent intensity than those that are. Each group had ten flies to make ten trials for each of the three control groups and the experimental group. Each fly counts as one trial, as the microscopy of the brains was performed.

**Figure 2:**
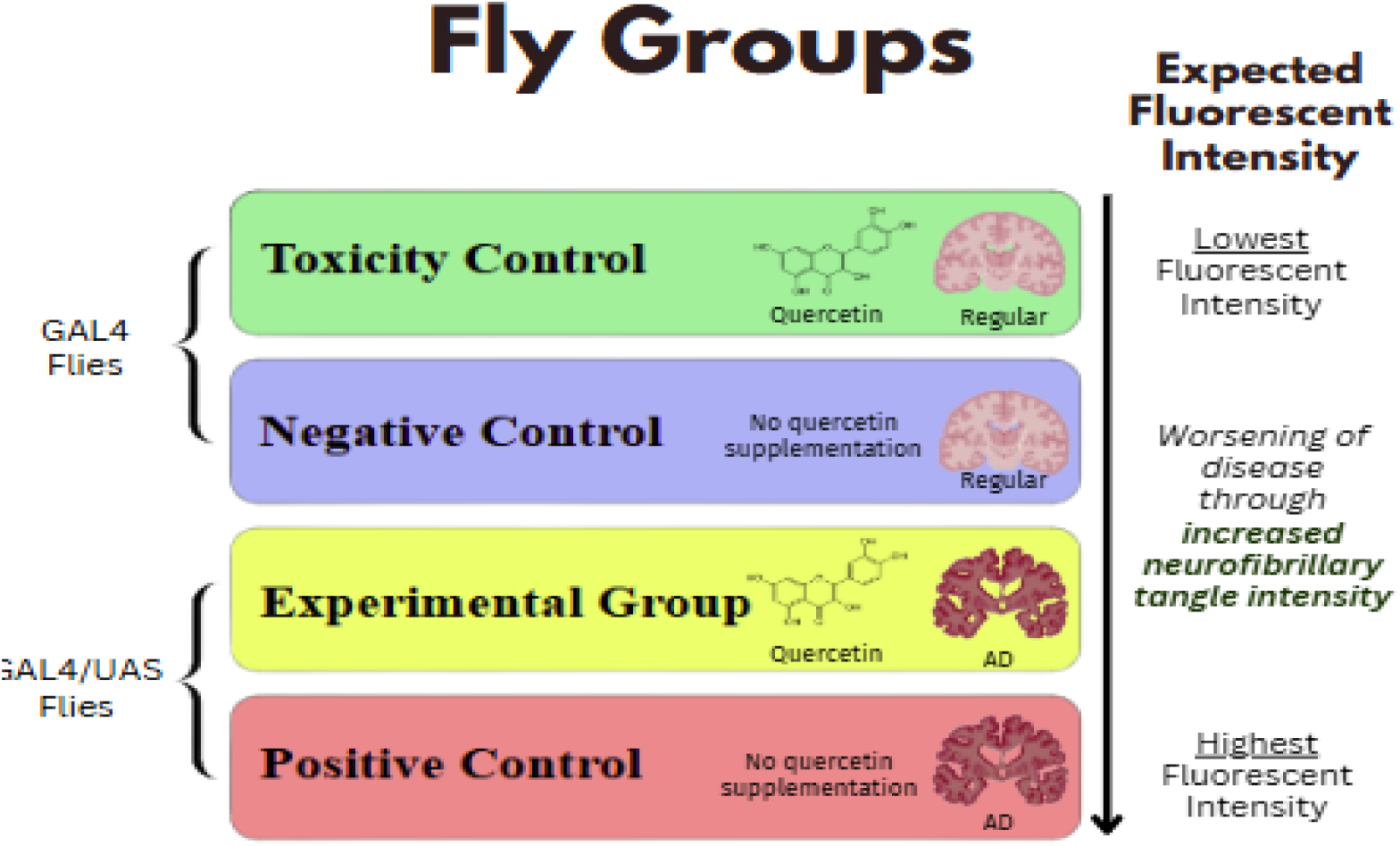
Groups of *Drosophila melanogaster* and expected results

## Materials

● UAS Line to express human tau in *Drosophila* (y[1] w[1118]; P{w[+mC]=UAS-MAPT.A}59A) (Bloomington *Drosophila* Stock Center #181) <u>BSL</u> <u>Level-1</u>
● GAL4 Line to express the proteins in the neurons and have a balancer chromosome (CyO) to indicate which progeny have both the GAL4-elav and UAS-MAPT chromosomes (P{w[+mC]=GAL4-elav.L}2/CyO) (Bloomington *Drosophila* Stock Center #8765) <u>BSL Level-1</u>
● Quercetin (AOS)
● (C.I. 14780) 112580-Thiazine Red (DC Fine Chemicals)
● Phosphate buffered saline (AOS)
● Lab equipment (AOS)

○ Freezers
○ Ice bath
○ Panasonic Genius Sensor NN-SN936W 1250 Watt Model Microwave
○ Leica Brightfield Fluorescence Microscope
○ Pipettes
○ Thermometer
○ Stirring rods
○ Beaker (1 L and 500 mL)
○ Graduated cylinders (500 mL and 100 mL)
○ Weighboats
○ Weigh scale
○ Weigh paper
○ Scoopula
○ Tweezers
● Carbon Dioxide (CO_2_) Sorting Materials (AOS)

○ CO_2_ tank
○ CO_2_ gun
○ CO_2_ pad
○ Labomed Luxeo 6z Microscope
○ Feather
○ Paintbrush
● *Drosophila melanogaster* food (AOS)

○ Distilled water
○ Cornmeal
○ Yeast
○ Agar
○ Soy flour
○ Light corn syrup
○ Propionic acid
○ Cheesecloth
● Fly storage materials

○ Medium size clear plastic vials
○ Foam plugs (flugs)
● Safety Supplies

○ Lab coat
○ Goggles
○ Close-toed shoes
○ Gloves
○ Hot hands

### Tapping Flies

First two vials were collected, a vial containing flies and an empty vial containing fly food. The flug of the vial containing flies was loosened and the flug of the vial containing the food was removed. The closed vial containing the flies were held in the left hand, while the open vial containing the fly food was held in the right hand. The vial containing the flies was then firmly tapped against a flat surface. As a result of this tapping, the flies fell to the bottom of the vial, and at this point, the flug of the fly vial was flicked off. The opening of the vial containing food was quickly placed over the vial containing flies, which created a seal so that the flies couldn’t escape. With the vials on top of each other, both vials were flipped upside down while maintaining the seal. Then, the vials were firmly tapped against the surface again to send flies to the bottom of the new food vial, and the flug was quickly inserted into the new vial while keeping the old vial upside down. Once the new vial was closed, the flug for the old vial was replaced. Finally, the new vial was labeled with the date, initials, and the type of fly stock. The old vial was also labeled again with the date that the tapping took place. Both vials were placed back into the tray and stored at room temperature.

### Maintaining Flies

Vials were tapped every four days to expand a stock, and stocks were maintained by tapping flies every three weeks. Moreover, if vials were dehydrated, a spray bottle with deionized water was used to spray onto the flug. Additionally, to prevent mites and mold, tools, utensils, and station was wiped with 70% ethanol before taking the flies out the vial to sort. If mites and mold were found, then the inside of the vial was sprayed with 70% ethanol and the vial was discarded in the large trash can.

### Fly Food Preparation

390 mL of distilled water was added into a beaker, and a weigh scale was used to measure 6.75 grams of yeast, 3.90 grams of soy flour, 28.50 grams of cornmeal, and 2.25 grams of agar into weight boats. Afterwards, 30 mL of corn syrup was measured using the scale into a 1 L beaker. The distilled water was combined into the beaker with the corn syrup, and the mixture was stirred using a stir rod to dissolve the corn syrup. Then, the yeast, sour flour, cornmeal, and agar amounts were added into the 1 L beaker, and the mixture was mixed again using a stir rod. If quercetin supplemented food was being made, the appropriate amount of quercetin would be added and stirred into this mixture as well. For 1 mM, 10 mM, and 100 mM concentrations, 0.14 grams, 1.4 grams, and 14 grams of quercetin were added, respectively. For a half batch of food, all ingredient values were halved and for a quarter batch, all values of ingredients were quartered. Once dry ingredients were added, the mixture was placed in the microwave and heated at 30 second intervals for either a half or quarter batch until the mixture bubbled and rised. If a full batch of food was being made, then 60 seconds intervals were used. After each interval, the mixture was removed using hot hands and was stirred. After the mixture boiled, the mixture was placed into the microwave until the mixture boiled two more times. Then, the food was left to cool to 70°C using a thermometer, and a cheese cloth was added over the beaker to cover the food in order to prevent contamination while cooling. The cheese cloth was sealed on the top of the beaker using a rubber band or a heavy object. Once the food reaches 70°C, 1.88 mL of propionic acid was added to the food using a pipette and mixed using a stir rod. This value was changed based on if the batch was a quarter or half batch of food. Then 10 mL of the resulting mixture was added into new vials, and the vials were covered with a cheesecloth. Once the food had cooled, flugs were inserted into the vials, and the vials were placed into the cardboard food tray. This tray with the food vials was placed in the refrigerator until future use.

### Toxicity Assay

For each concentration (1 mM, 10 mM, and 100 mM), ten female GAL4 flies were fed the concentration of quercetin into their daily diet for a two week period. 0.14 grams, 1.4 grams, and 14 grams of quercetin were needed to make 1 mM, 10 mM, and 100 mM quercetin concentrations. Quercetin amounts are calculated based on concentration, and the amounts will be added to the fly food mixture. The molar mass of quercetin is 302.236 g/mol, and the amount of fly food made is 463.28 mL, which was calculated based on the fly food preparation amounts (390 mL + 30 mL+ 6.75 g + 3.9 g + 28.5 g + 2.25 g + 1.88 mL). A sample calculation of determining the grams of quercetin needed for the 1 mM concentration is shown as below. The process in calculating quercetin amounts were repeated for 10 mM and 100 mM concentrations.

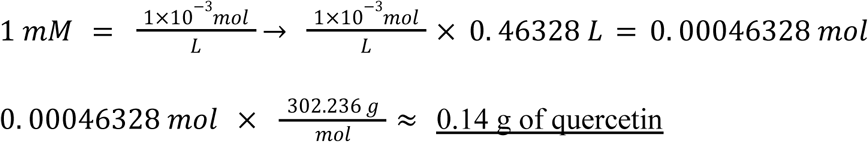

The amount of flies each day were recorded in order to determine the maximum amount of quercetin that would be fatal and the optimal quercetin to test was chosen based upon the results. The concentrations with the highest survival rates were used for experimentation.

### Fly Sorting

To sort flies into male and female, the genitalia of flies were inspected. While there are some characteristics that determine male versus female flies, the only reliable indicator of sex are the genitalia. Males contain genetial claspers with a blunt end, are usually smaller, have darker and rounder abdomens, and contain knee pads on the front legs, called sex combs. Female flies contain pointy ovipositor ends, are usually bigger, and have stripes down their body. Furthermore, virgin flies were sorted by looking for a meconium patch and folded wings. Virgin flies are also usually paler and larger than non-virgin flies. Once flies were found to be virgin, the gender of the fly was determined based on the genitalia described above.

### CO_2_ Sorting

CO_2_ sorting was performed to distinguish the gender of flies in order to make crosses and to find female flies for experimentation. Firstly, the CO_2_ wheel was turned clockwise to turn the CO_2_ tank on, and both a CO_2_ gun and a CO_2_ box are attached to the tank. The CO_2_ coming out was at a low level, and the sound of the gas was barely heard. The closed vial was held sideways, and the CO_2_ gun was placed inside the vial by slipping the needle of the gun through the side of the flug. Once the flies passed out, the flug was removed and flies were placed on the CO_2_ box under the Labomed Luxeo 6z Microscope. The overhead light of the microscope was turned on, and magnification/focus was adjusted as needed. The top wheels controlled the magnification, while the side wheels controlled the focus. Using a feather, flies were sorting through based on the gendering/virgining protocol. The sorted flies were then placed in different vials, and the flies were left sideways until they woke up. Males, females, virgin males, and virgin females were all kept in separate vials.

### Genetic Cross

Firstly, virgining and gendering protocols were followed to find GAL4 virgin females and UAS males. Six GAL4 virgin females were placed into the same vial as two UAS males. Once the male flies and virgin females mated, the flies were tapped into a new vial. This resulted in a vial with the eggs, and the new vial with the cross was dated and labeled. Around two weeks later, the progeny from the mating of the GAL4 and UAS flies hatched, and these progeny followed the CO_2_ sorting protocol to be placed under the Labomed Luxeo 6z Microscope. The progeny were examined, and the flies of the desired alleles were distinguished to find the progeny that expressed Alzheimer’s disease. Progeny with curly wings were removed from the experimental Alzheimer’s disease flies that contained straight wings. From the experimental flies, female flies that expressed Alzheimer’s disease were found by following the gendering protocol. The female GAL4/UAS flies that expressed Alzheimer’s disease were placed into a new vial, and the vial was placed sideways until the flies woke up. Then, the new vial was dated and labelled, and the vial containing the eggs of the crossed flies were discarded. Additionally, the flies of the cross that do not exhibit Alzheimer’s disease were discarded as well. Flies were discarded by either placing them into the -20°C freezer for one hour, or by spraying ethanol into the vial and then disposing the vial with the flug on.

Flies with elav-GAL4/CyO and UAS-MAPT were crossed to produce progeny exhibiting Alzheimer’s disease. The parent genotypes were yw; UAS-MAPT and w^-^; elav-GAL4/ CyO. The elav-GAL4 and CyO were the male *Drosophila melanogaster* genes and the UAS-MAPT will be the female *Drosophila melanogaster* genes. Progeny containing elav-GAL4 and UAS-MAPT will exhibit Alzheimer’s disease and these did not have curly wings. These were the progeny that were desired for the use of experimentation as they exhibited Alzheimer’s disease. Progeny containing the UAS-MAPT and CyO genes will have curly wings.

### Brain Dissection (Method adapted from Kelly et al. 2017)

Once five weeks old and ready to be dissected, flies were killed through ice. Tweezers were used to place one fly on the Labomed Luxeo 6z Microscope microscope into 1 mL of 0.01 M phosphate buffered saline (PBS) in the center of a dissection dish. Pins were placed into the abdomen and chest of the fly, and then the mouth and head were taken off using two tweezers. Each tweezer was placed on one eye, and the tweezers were pulled away from the fly to rip each eye apart. Afterwards, the connecting part of the eyes and the section covering the brain was carefully taken away with the tweezers. Then, the brain of the fly should remain, and the brain should look translucent, jelly-like, and sticky. If data was not immediately able to be collected, then the brain was frozen at -80 degrees Celsius.

### Brain Staining (Method adapted from Calvo-Rodriguez et al., 2019 and Luna-Muñoz et al., 2008)

Firstly, if the brain was frozen, the brain was left to thaw for approximately 15 minutes. Brains were washed with phosphate buffered saline for 5 minutes, and then the brains were stained for 20 minutes with 1.6 μg/ml of the Thiazine Red dye. Afterwards, brains were taken out of the phosphate buffered saline, and they were placed in another dish for quantification of the neurofibrillary tangles under the fluorescent microscope.

### Quantifying Tangles (Method adapted from Kniewallner et al., 2016)

The Leica Brightfield Fluorescence microscope and the computer connected to the microscope were turned on. Leica Application Suite X software on the connected computer was opened and started up. The microscope and digital camera were checked to be both plugged into the power strip along the table, and the green button labeled “shutter” on the control pad was clicked so that the laser glowed green and highlighted the fluorescent parts of the brain. Additionally, the light intensity knob was turned down to the lowest setting by turning counterclockwise. The black turret above the objectives was set to 2, and this setting was labelled “GFP”. The silver knob attached to the microscope was pulled out to connect to the digital camera on the software. Microscope slides for the brains were then created by following a wet mount technique using phosphate buffered saline. A drop of phosphate buffered saline was placed on the slide and the brain was inserted onto the saline using tweezers. A coverslip was placed over the brain and the phosphate buffered saline, and the slide was placed underneath the microscope. The focus and magnification of the microscope was adjusted as needed, and the image of the brain would be displayed on the computer as well. The pause button on the bottom-left corner of the software was clicked to pause the live image, and the image was right clicked to showcase “Export”. The image would then be saved into a folder stored on the computer. Once imaging was done, these images were sent and saved through Google Drive in order for quantification to occur.

### Image J Protocol

The brain image was first imported into the Image J software, and the cursor was dragged over an area of the image to measure the average fluorescence intensity in that area. The region measured was a constant 35 *u*^2^ and the intensity was measured in the middle of the brains for all trials. The key “t” on the keyboard was clicked to add a region to the ROI (region of interest) manager. “Analyze” on the top row was selected, and from the dropdown, “Measure” was clicked. Once “Measure” was clicked, the size of the area, the average fluorescence intensity, and the skew was showcased. The mean fluorescent intensity was then recorded into the data table, and this process was repeated for all measured brain images. A breakdown of the methods will be shown below in Figure 3.

**Figure 3:**
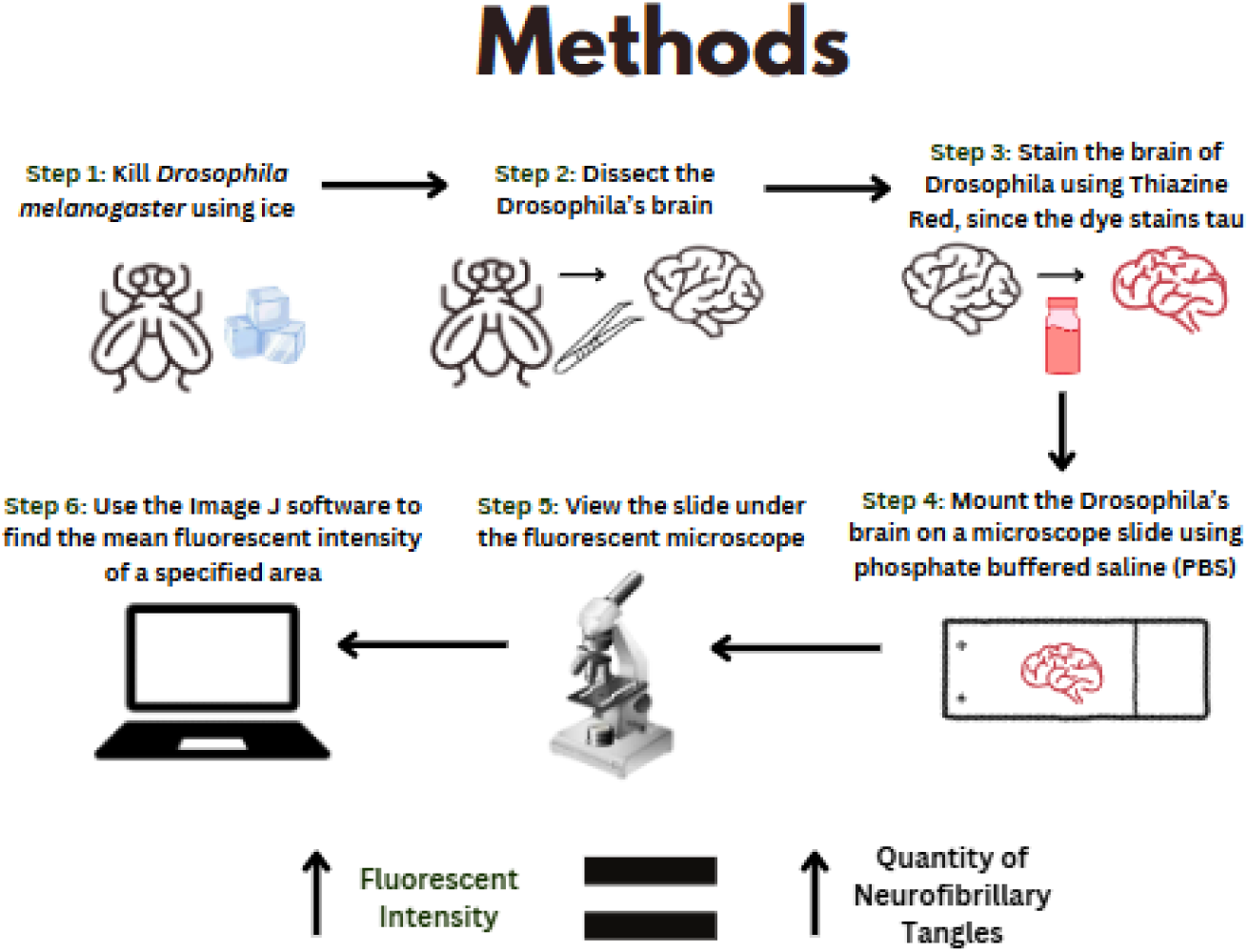
Visual representation of methods

### Safety

1. Safety data sheets

a. Thiazine Red SDS
b. Quercetin SDS
c. Propionic Acid SDS
d. Phosphate Buffered Saline SDS

The general safety procedures performed were wearing closed-toed shoes in the lab and during experimentation, wearing goggles and a lab coat when handling chemicals and glassware, and wearing gloves when handling the chemicals. Moreover, heat resistant mitts were worn when the microwave was used to make food in order to prevent burns. Flies were safely disposed of through freezing or ethanol, and hands were washed routinely after experimentation. If any of the chemicals were inhaled, then the person was moved to fresh air. If chemicals were ingested, then the mouth was rinsed with water. Additionally, eye contact with the chemicals was avoided because the chemicals used could cause serious eye irritation.

In regards to chemical disposal, pipette tips used to measure propionic acid and phosphate buffered saline were disposed of in a special waste disposal bag. Quercetin, Thiazine Red, propionic acid, and phosphate buffered saline was also collected into a chemical-safe, labeled, and sealed container for disposal. Thiazine Red, quercetin, propionic acid, and phosphate buffered saline was disposed of into an approved waste disposal plant.

Quercetin was refrigerated and moved away from heat, moisture, and direct sunlight. Thiazine Red was stored in a well ventilated area at room temperature and away from sources of ignition. Propionic acid was stored in a well ventilated area, kept cool, and placed to avoid heat, sparks, and open flames. Phosphate buffered saline was stored in a well ventilated area and was kept cool as well.

In the event that quercetin was spilled, all sources of ignition would be removed, and a solid spill material with 70% ethanol would be used to pick up the spill. The absorbent would be sealed into a plastic bag for disposal. The corresponding chemical disposal protocol would be followed. Additionally, the contaminated surfaces would be washed with 70% ethanol and would be followed by a soap and water solution. In the event that Thiazine Red, propionic acid, or phosphate buffered saline was spilled, the spill would be absorbed with inert material. Then, the inert material would be placed into a suitable disposal container. The corresponding chemical disposal protocol would be followed.

In the event of skin contact with chemicals, the chemical would be quickly washed off using the sink or the safety shower in severe cases. An adult would be notified during severe cases. In the event of contact with chemicals in the eyes, an individual would be moved to the eye wash station, and the eyewash station would be used for 15 minutes. If applicable, contact lenses would be removed. Additionally, an adult would be notified in this case as well. Lastly, in the event of broken glass, an adult would be notified to clean up the glass. A broom and dustpan would be used and broken glass would be placed into the glass disposal cardboard box.

## Data

**Table 1:** Toxicity Assay-Number of Flies Alive at 1, 10, & 100 mM Quercetin Concentrations.

| Days after start | Number of Flies Alive |  |  |
| --- | --- | --- | --- |
|  | Concentration |  |  |
|  | 1 mM | 10 mM | 100 mM |
| 0 | 10 | 10 | 10 |
| 1 | 10 | 10 | 6 |
| 2 | 10 | 10 | 6 |
| 3 | 10 | 10 | 6 |
| 4 | 10 | 10 | 6 |
| 5 | 9 | 10 | 6 |
| 6 | 9 | 10 | 6 |
| 7 | 9 | 10 | 6 |
| 8 | 9 | 10 | 6 |
| 9 | 9 | 10 | 6 |
| 10 | 9 | 10 | 6 |
| 11 | 9 | 10 | 6 |
| 12 | 9 | 10 | 6 |
| 13 | 9 | 10 | 6 |

**Table 2:**
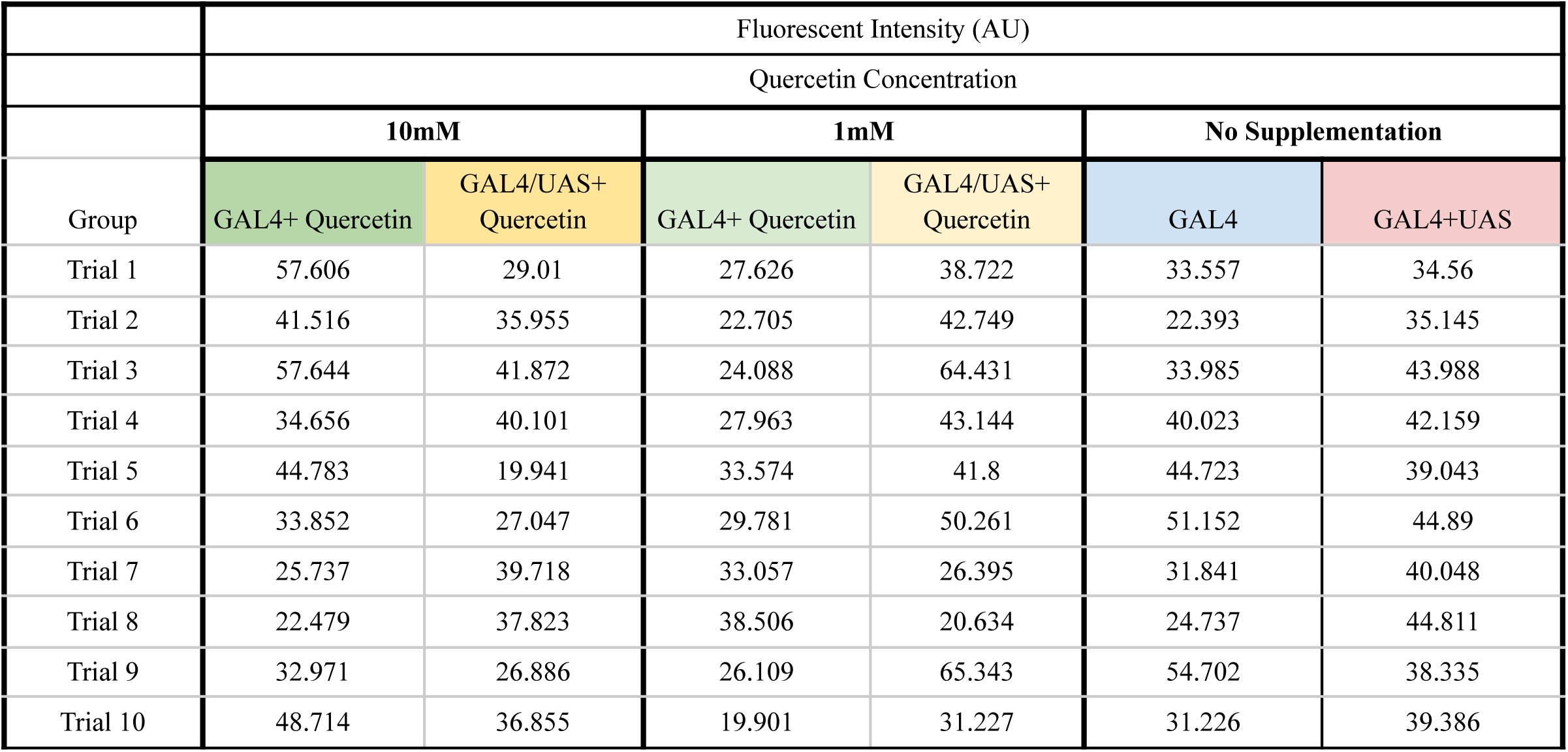
The Relationship Between Fly Groups and the Fluorescent Intensity (AU) of Neurofibrillary Tangles.

### Results

**Table 3:** Means Fluorescent Intensity Values of All Fly Groups.

|  | Fluorescent Intensity (AU) |  |  |  |  |  |
| --- | --- | --- | --- | --- | --- | --- |
|  | Quercetin Concentration |  |  |  |  |  |
|  | 10mM |  | 1mM |  | No Supplementation |  |
| Group | GAL4+ Quercetin | GAL4/UAS+ Quercetin | GAL4+ Quercetin | GAL4/UAS+ Quercetin | GAL4 | GAL4+UAS |
| Mean | 40.317 | 38.162 | 28.331 | 49.609 | 40.237 | 36.834 |

**Table 4:** ANOVA Statistical Analysis on Fly Groups and the Fluorescent Intensity (AU) of Neurofibrillary Tangles.

| **Effect** | **Degrees of Freedom**<br>(df) | **F-statistic**<br>(MS / MS residual) | **P-value** | **Interpretation of P** |
| --- | --- | --- | --- | --- |
| Group Of Fly | 5 | 2.8 | 0.02 | A P-value of 0.02 means strong confidence that the groups are different. |
| Error or Residual | 54 | NA | NA | NA |

**Table 5:** Post-Hoc Tukey’s Range Statistical Test on Fly Groups and the Fluorescent Intensity (AU) of Neurofibrillary Tangles.

| **Pair** | **Mean Diff** | **P-value** | **Interpretation of P-value** |
| --- | --- | --- | --- |
| GAL4/UAS10mM - GAL4/10mM | -6.47 | 0.68 | Means: no evidence that the groups might be different |
| GAL4/1mm - GAL4/10mM | -11.7 | 0.10 | Means: some indication that the groups might be different |
| GAL4/UAS1mM - GAL4/10mM | 2.47 | 0.99 | Means: that the groups are NOT different |
| GAL4 - GAL4/10mM | -3.16 | 0.98 | Means: no evidence that the groups might be different |
| GAL4/UAS - GAL4/10mM | 0.241 | 1.00 | Means: that the groups are NOT different |
| GAL4/1mm - GAL4/UAS10mM | -5.19 | 0.84 | Means: no evidence that the groups might be different |
| GAL4/UAS1mM - GAL4/UAS10mM | 8.95 | 0.33 | Means: little to no indication that the groups might be different |
| GAL4 - GAL4/UAS10mM | 3.31 | 0.97 | Means: no evidence that the groups might be different |
| GAL4/UAS - GAL4/UAS10mM | 6.72 | 0.65 | Means: no evidence that the groups might be different |
| GAL4/UAS1mM - GAL4/1mM | 14.1 | 0.03 | Means: strong confidence that the groups are different |
| GAL4 - GAL4/1mM | 8.5 | 0.39 | Means: little to no indication that the groups might be different |
| GAL4/UAS - GAL4/1mM | 11.9 | 0.09 | Means: some confidence that the groups are different |
| GAL4 - GAL4/UAS1mM | -5.64 | 0.79 | Means: no evidence that the groups might be different |
| GAL4/UAS - GAL4/UAS 1mM | -2.23 | 1.00 | Means: that the groups are NOT different |
| GAL4/UAS - GAL4 | 3.4 | 0.97 | Means: no evidence that the groups might be different |

**Graph 1:**
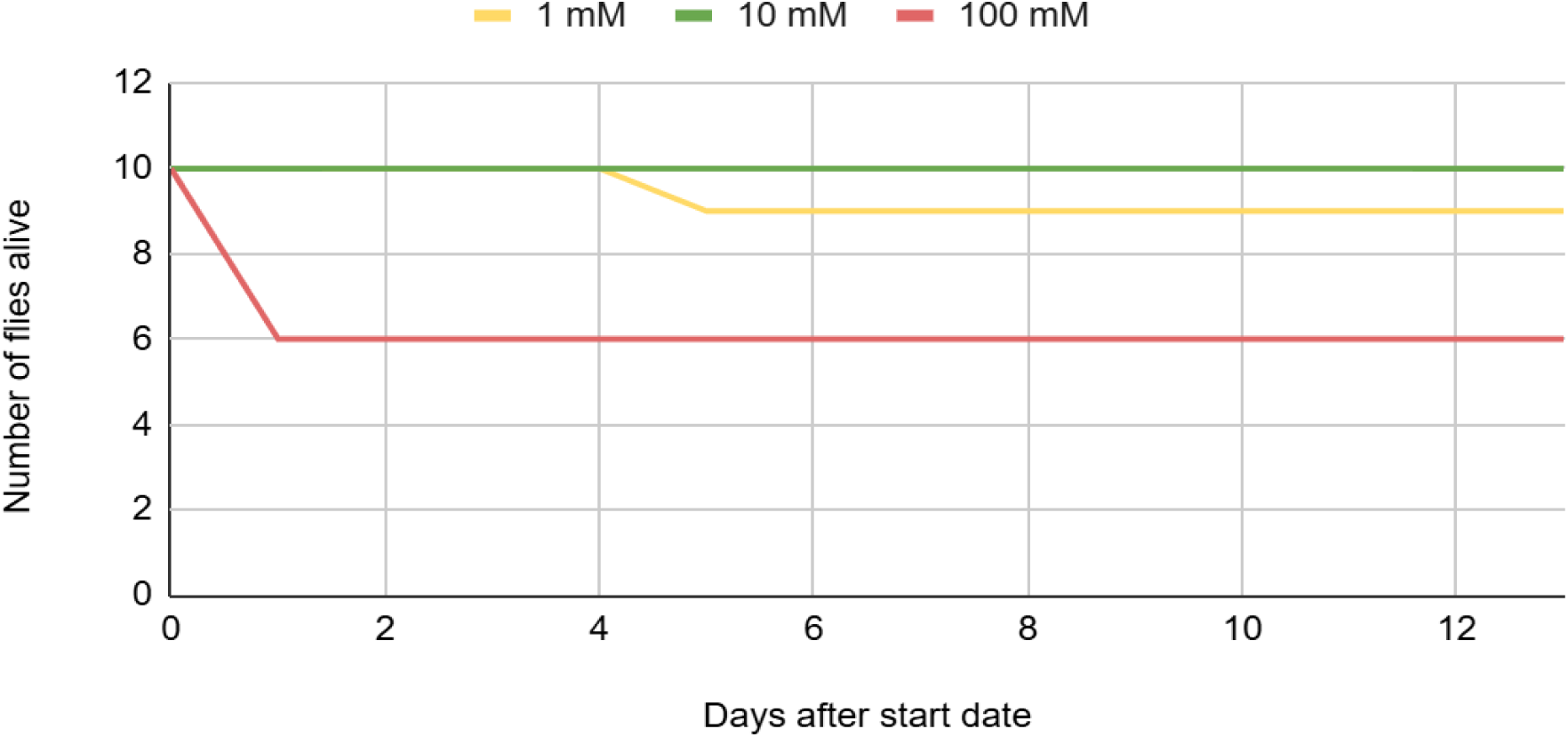
Days Elapsed Over Two Weeks vs. the Number of *Drosophila melanogaster* Alive

**Graph 2:**
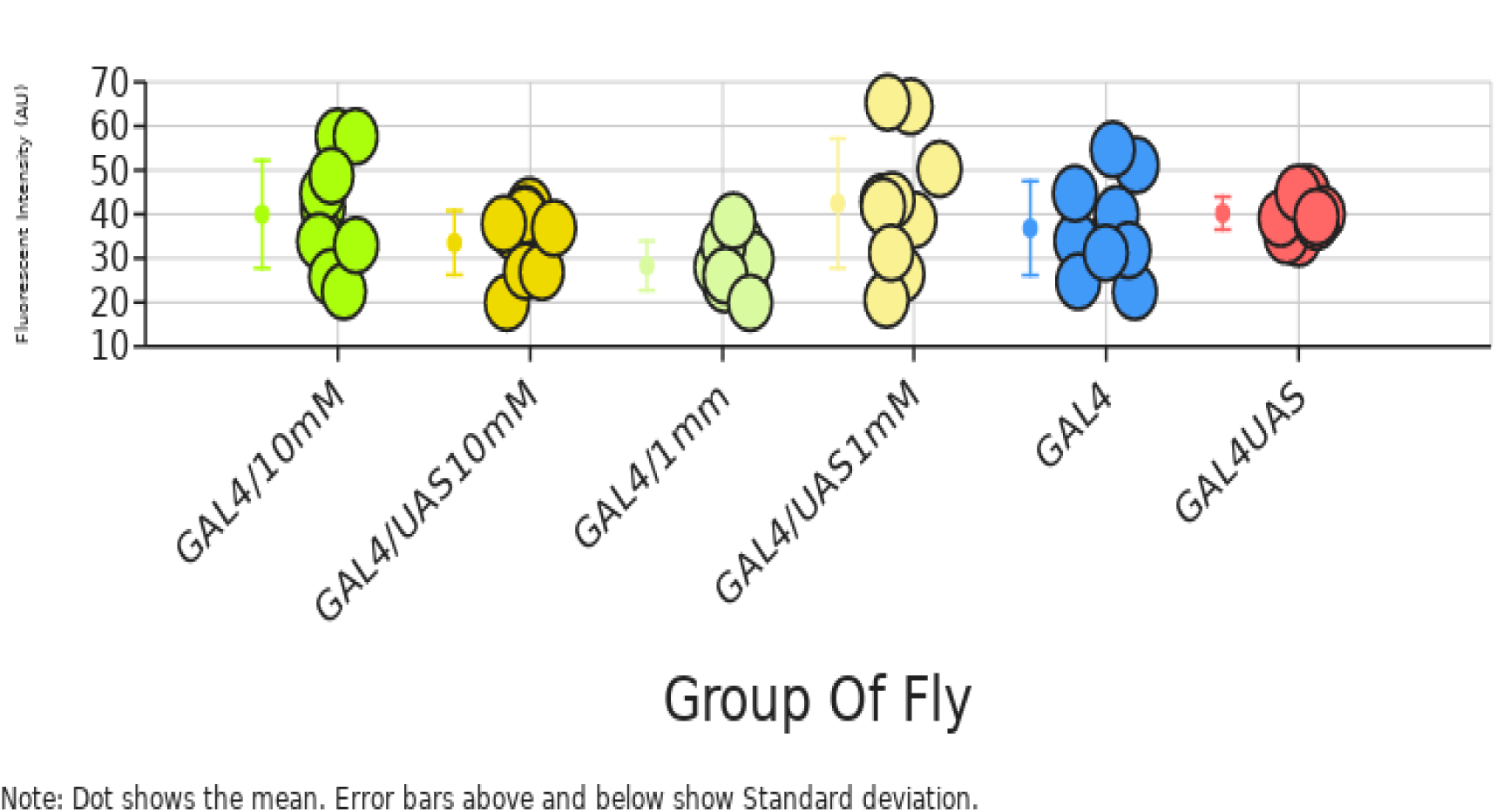
Fluorescent Intensity (AU) of Fly Brains Based On Fly Groups

For this project, the quantity and fluorescence intensity of neurofibrillary tangles will be tested and compared. There is a negative control, a toxicity control, a positive control, and one experimental group. There are ten flies per group to make ten trials. However, time can be a limiting factor for the number of trials. This is because it takes time for flies to be shipped and because 5 weeks have to pass for Alzheimer’s disease to be exhibited in the flies. Additionally, dissecting brains takes a long amount of time so only 10 brains were done per group. Moreover, all the trials are limited to be done within the scope of one year. The ANOVA and Post-hoc Tukey test was used to determine if there are statistically significant differences between different groups.

## Discussion

The concentrations of 10 mM and 1 mM were chosen as the most optimal concentrations to test as the least amount of flies had died over the two week toxicity testing period from these concentrations. By the end of the two weeks, no flies died in the 10 mM concentration and only one fly died in the 1 mM concentration. This indicated that 10 mM was the optimal concentration as there were no adverse side effects and the 1 mM was found to be the second most optimal concentration of quercetin because there was only one death. The data suggests that quercetin supplementation has no significant impact on the intensity of the neurofibrillary tangles within the brains of the *Drosophila melanogaster* that exhibited Alzheimer’s disease. There was no significant difference between GAL4/UAS and GAL4/UAS + quercetin groups, as the p-values were above 0.05 as the p-value for the difference between GAL4/UAS GAL4/UAS10mM was 0.67 and the p-value for the difference of GAL4/UAS - GAL4/UAS 1mM was 1.00. These extremely high p-values were a clear indication that quercetin did not have a statistically significant effect on the fluorescent intensity of the neurofibrillary tangles.Furthermore, quercetin supplementation had no statistical impact on the neurofibrillary tangles within *Drosophila* that did not have Alzheimer’s disease. Our results thereby indicate that quercetin supplementation does not have a positive nor negative impact on neurofibrillary tangles within *Drosophila melanogaster* which was not as hypothesized, as we had previously believed that quercetin supplementation would have a positive effect on neurofibrillary tangles because quercetin is a strong antioxidant and could potentially have potent binding effects to prevent tau from misfolding and turning into toxic aggregates of neurofibrillary tangles.

Since the results indicate that quercetin supplementation has no significant effect on the intensity of the neurofibrillary tangles, quercetin would not be effective as an oral supplement used to prevent Alzheimer’s disease as quercetin supplementation would not have a significant impact on one of the main mechanisms of the Alzheimer’s pathology. However, since our data is prone to a lot of error and there was significant error present within our data collected, we cannot forgo the idea that quercetin could have a positive impact on the symptoms of Alzheimer’s disease. There was a lot of variation within our data which showed that our method of data collection was not very precise. Additionally, not much data was collected as only ten fly brains were analyzed for each fly group. As smaller sample sizes are more susceptible to error, there is a chance that our results are flawed. Issues in our Alzheimer’s disease model itself were also showcased through the Post-Hoc test, as the p-value between the GAL4 and GAL4/UAS group was not significant because the p-value was above 0.05. Moreover, quercetin could have an impact on other mechanisms of Alzheimer’s disease such as amyloid beta plaques and oxidative stress, especially since quercetin is a potent antioxidant that could reduce oxidative stress.

The main source of error within this experiment was the fluorescence of the brain tissue and hair fibers that were not removed. This could have drastically skewed the results by bringing up the mean fluorescence of each brain due to this extra fluorescence of the part of the brain which was not being studied. Also, there was a large central area of 35 u^2^ which was measured which could have skewed the data because there was a larger chance of this extra brain matter and hair being part of the brain which was being imaged and analyzed. Additionally, there was a chance that the Thiazine Red had bound to tau in general, instead of the neurofibrillary tangle aggregates that were trying to be studied. As there is healthy tau present in the brain which is necessary for brain function, if the fluorescent intensity measured the amount of tau within the brain, neither a positive nor negative effect could be concluded as the researchers would not have known if this tau had misfolded and turned toxic or if the tau was healthy for the brain. Additionally, as there was varied waiting times between staining, dissecting, and analyzing the brain under the microscope, the brains could have been exposed to the dye for longer periods of time which would have brought up the mean fluorescent intensity or the tangles could have dissipated due to an amount of time passing after the death of the *Drosophila melanogaster*.

## Conclusion

In general, the data indicated that the supplementation of quercetin within the food of flies did not have a significant impact on the intensity of neurofibrillary tangles, as the p-values were above 0.05. Neither the *Drosophila* that expressed Alzheimer’s disease nor those that did not express Alzheimer’s disease had not shown any strong indication of the improvement of the intensity of the neurofibrillary tangles within their brains. Our data collection had many errors present so some of the data could be due to these errors, however, the original data collected cannot be overlooked and the data makes it seems like there is no benefit for patients with Alzheimer’s disease to take quercetin as an oral supplement or people to take quercetin to prevent themselves from getting Alzheimer’s disease. Other symptoms of Alzheimer’s disease could be tested such as memory or the strength of the amyloid beta plaques, however, these were not tested this year.

## Future Work

In the future, quercetin could be tested on the short term and long term memory loss of *Drosophila melanogaster* exhibiting Alzheimer’s disease through the use of the aversive phototaxic suppression assay. This assay would indicate if the quercetin supplementation had an effect on the memory loss associated with Alzheimer’s disease through the use of different smells. In this assay, the flies are trained to associate light, which they have a natural preference towards, with a bad smell. They are continuously tapped between vials and learn that the dark vial is better for them. Then, after a waiting period, the flies are placed into two connected vials, one dark and one light, and the number in each are counted. If more flies go into the dark vial, then they have learned that the light vial is bad due to the smell and therefore, they have better memory due to them remembering the difference after the training period. The effect of quercetin supplementation could easily be analyzed by having a group of flies with Alzheimer’s that was supplemented with quercetin and one which was not supplemented with quercetin undergo this assay. If more flies went to the dark vial which took quercetin, then the quercetin reduced the memory loss effect of Alzheimer’ s disease. Additionally, more research could be done on antioxidants in general as there has not been much research on their therapeutic effects as supplements within patients who exhibit Alzheimer’s disease. As antioxidants are able to bring down the oxidative stress levels within the brain, they could have a significant impact on neurodegeneration and memory loss within Alzheimer’s. Furthermore, more research is needed on compounds that bind tau as binding tau is the most effective way to prevent the formation of neurofibrillary tangles. If more compounds are identified that are able to bind with tau and prevent fibril formation, then a similarity could be found between these compounds which could provide insight into possible ways to prevent the exhibition of Alzheimer disease and slow down the progression of the disease within patients.

